# Painting the evolutionary history of a multicompartmentalized syngameon in North American *Castilleja* (Orobanchaceae)

**DOI:** 10.64898/2026.09.08.750233

**Authors:** Malia A. Santos, Sarah J. Jacobs, Maribeth Latvis, Sean Harrington, David C. Tank

## Abstract

Understanding the evolutionary history of recent rapidly radiating groups remains a major challenge in systematics, especially when incomplete reproductive isolation and rampant interspecific gene flow obscure phylogenetic signal. Here, we present a comprehensive phylogenomic study of the genus *Castilleja* (“the paintbrushes”, Orobanchaceae), integrating nuclear and chloroplast data to investigate patterns of diversification, hybridization, and geographic structure across its range. Our results provide the first evidence that *Castilleja* functions as a multicompartmentalized syngameon—a network of closely related species with ongoing or historical gene flow among non-sister lineages within and between geographically structured compartments. High nuclear gene tree discordance, coupled with strong cpDNA geographic compartment structuring and nuclear network analysis, supports the hypothesis that recent rapid radiation and Plio-Pleistocene climatic fluctuations created alternating periods of range expansion and contraction. These dynamics likely promoted localized hybridization within isolated compartments during contraction and further introgression between compartments during range expansion. Our findings not only clarify the evolutionary relationships within *Castilleja* but also demonstrate the importance of genome-scale data and syngameon theory in understanding diversification in lineages where hybridization is rampant.

**Significance Statement:** Groups in which non-sister species frequently hybridize and exchange genes, yet maintain distinct identities, form what is known as a syngameon. Using a phylogenomic dataset, we show that the genus *Castilleja* exhibits extensive historical and contemporary gene flow across numerous species and broad geographic regions, producing a complex evolutionary history best described as a multicompartmentalized syngameon. Unlike most documented syngameons, which involve relatively few interacting species within the same geographic area, *Castilleja* consists of multiple geographically and ecologically structured compartments connected through repeated cycles of range expansion, isolation, and secondary contact during the Plio-Pleistocene. Our findings suggest that multicompartmentalized syngameons may represent a common but underrecognized outcome of rapid radiations.

## Introduction

Hybridization and introgression have most often been studied in sister species, but many studies have now shown that reticulate evolution between non-sister lineages is much more common than once thought (e.g. oaks, pinyon pine, cichlids, Darwin’s finches) (1–8). In some clades, rampant interspecific hybridization makes it difficult to accurately infer species’ evolutionary histories using traditional phylogenetic approaches. This pattern is evident across the tree of life, and is particularly pervasive in plants, where in groups like oaks (*Quercus*)*, Iris,* birches (*Betula*), and willows (*Salix*), distinct morphological lineages can persist despite documented ongoing and persistent hybridization (4, 9–11). These taxonomic misfits have puzzled systematists for years, prompting the use of ‘syngameons’ to describe groups of organisms in nature that experience continuous or sporadic interspecies gene flow—estimated to occur in 25% of plants and 10% of animals (7, 12, 13). The concept of species connected by gene flow has been described multiple times to explain patterns observed in taxonomically complex groups. For example, Van Valen introduced the idea of a multispecies, or ‘a set of broadly sympatric species that exchange genes’ to explain these patterns in oaks (14). However, nearly 70 years earlier, Lotsy first articulated the syngameon concept to describe patterns seen in birches, where a group of closely related taxa form a complex network of hybrid combinations (15). Since these early descriptions, taxonomists have struggled to accurately depict the evolutionary history of such groups due to the limitations of morphological and single-locus data. Only with the advent of reduced representation genomic approaches—enabling efficient and cost-effective sampling across the genome—have we begun to unravel the complexity of these reticulate evolutionary patterns. Since then, many different hypotheses have arisen from the syngameon framework to explain the formation and persistence of these complex networks (13).

The rapid radiation hypothesis suggests that syngameons arise during rapid diversification, where short evolutionary timescales limit the development of complete barriers to reproductive isolation, thereby facilitating interspecific gene flow. In contrast, the edge-range hypothesis posits that the peripheral zones of species ranges are not a population sink; instead, when populations at range margins come into contact with different species, hybridization can occur, promoting persistence through introgression of beneficial alleles (13, 16). Both hypotheses highlight how incomplete reproductive barriers and dynamic geographic distributions can contribute to the structure of syngameons. While proposed as distinct hypotheses, we highlight that both can be acting concurrently to further complicate the history of a group.

Along with the confounding effects of interspecific hybridization in these recent radiations, incomplete lineage sorting (ILS) and paralogy resulting from gene duplication events and/or polyploidy further complicate efforts to infer bifurcating species trees. While traditional phylogenetic methods assume clear species boundaries and bifurcating evolutionary relationships, these assumptions often fail in the context of syngameons. To address this, bifurcating phylogenetic approaches paired with coalescent-aware phylogenetic methods and phylogenetic network analyses can more effectively model reticulate evolutionary histories while accounting for coalescent stochasticity. We use these techniques to quantify the amount of gene-tree discordance potentially attributable to syngameon processes in the taxonomically complex genus *Castilleja* (Orobanchaceae).

*Castilleja* is a rapidly radiating genus of ∼200 hemiparasitic species and, as we formalize here, represents a model case of a syngameon (Figure 1). It spans a wide latitudinal gradient distributed from Andean South America to Beringia (Figure 2E), with the majority of species diversity centered in Mexico and western North America, and occurs in a wide variety of habitats, ranging from mediterranean grasslands to alpine tundra. Interspecific hybridization in *Castilleja* is common and well documented, and both field and greenhouse studies have demonstrated that hybridization occurs freely between species, sometimes between different ploidy levels, and between species that are assumed to be distantly related based on morphological hypotheses of relationships in the clade (Figure 2B) (17–19). The degree and frequency of hybridization has undoubtedly led to the confounded and unresolved phylogenetic history within this clade (18, 20, 21). In addition to hybridization, the large majority of species are part of a perennial clade that is derived from a relatively species-poor grade of annual *Castilleja* species (20); this transition from an annual to a perennial lifestyle has been suggested to be a driver of diversification in the clade, where perennial species diversify at a ∼10X greater rate (20, 22, 23). Finally, polyploidy is rampant across perennial *Castilleja* species, where it is estimated that at least 50% of perennial species in western North America have at least some polyploid populations (20, 24–26).

**Figure 1.**
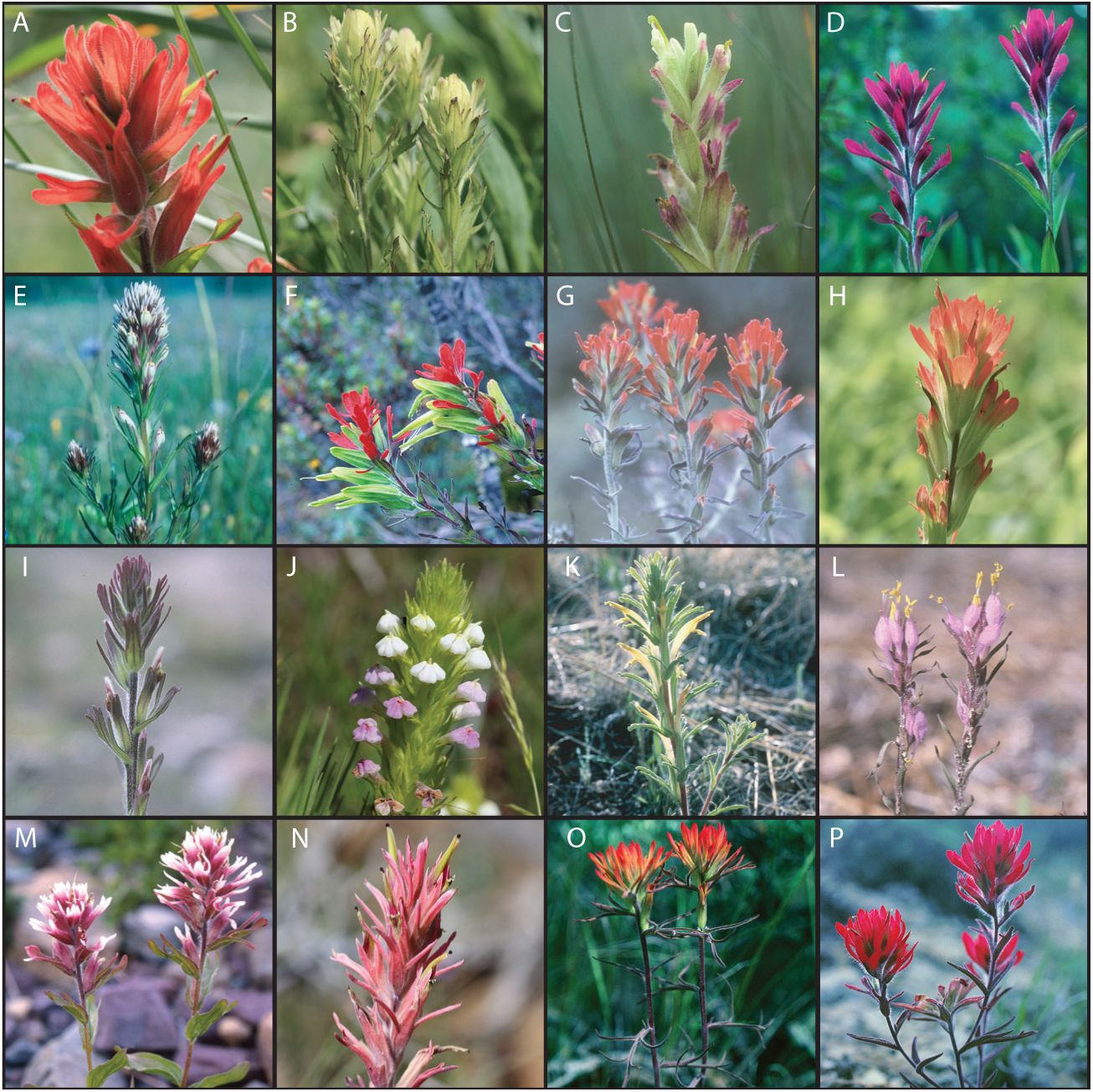
Photos of *Castilleja* species highlighting key morphological species groups that span the diversity seen in the group with name of morphological species group in parentheses (A-L): A) *Castilleja miniata var dixonii* (Miniata), B) *Castilleja cusickii* (Cusickii), C) *Castilleja chrysantha* (Cusickii), D) *Castilleja rhexiifolia* (Septentrionalis), E) *Castilleja densiflora* (Oncorhynchus), F) *Castilleja fissifolia* (Linariifoliae), G) *Castilleja foliolosa* (Lanatae), H) *Castilleja hispada var. acuta* (Chromosa), I) *Castilleja cerroana* (Cerroana), J) *Castilleja rubicundula* (Oncorhynchus), K) *Castilleja mexicana* (Callichroma), L) *Castilleja ophiocephala* (Ophiocephala). Putative hybrids observed wild with name of morphological species group in parentheses (M-P): M) *Castilleja rhexiifolia* (Septentrionalis) *x Castilleja occidentalis* (Septentrionalis), N) *Castilleja linariifolia* (Linariifoliae) *x Castilleja nana* (Pilosae), O) *Castilleja flava var rustica* (Callichroma) *x Castilleja chromosa* (Chromosa), P) *Castilleja scabrida var. barbeyana* (Chromosa) *x Castilleja chromosa* (Chromosa). All images © Mark Egger.

**Figure 2.**
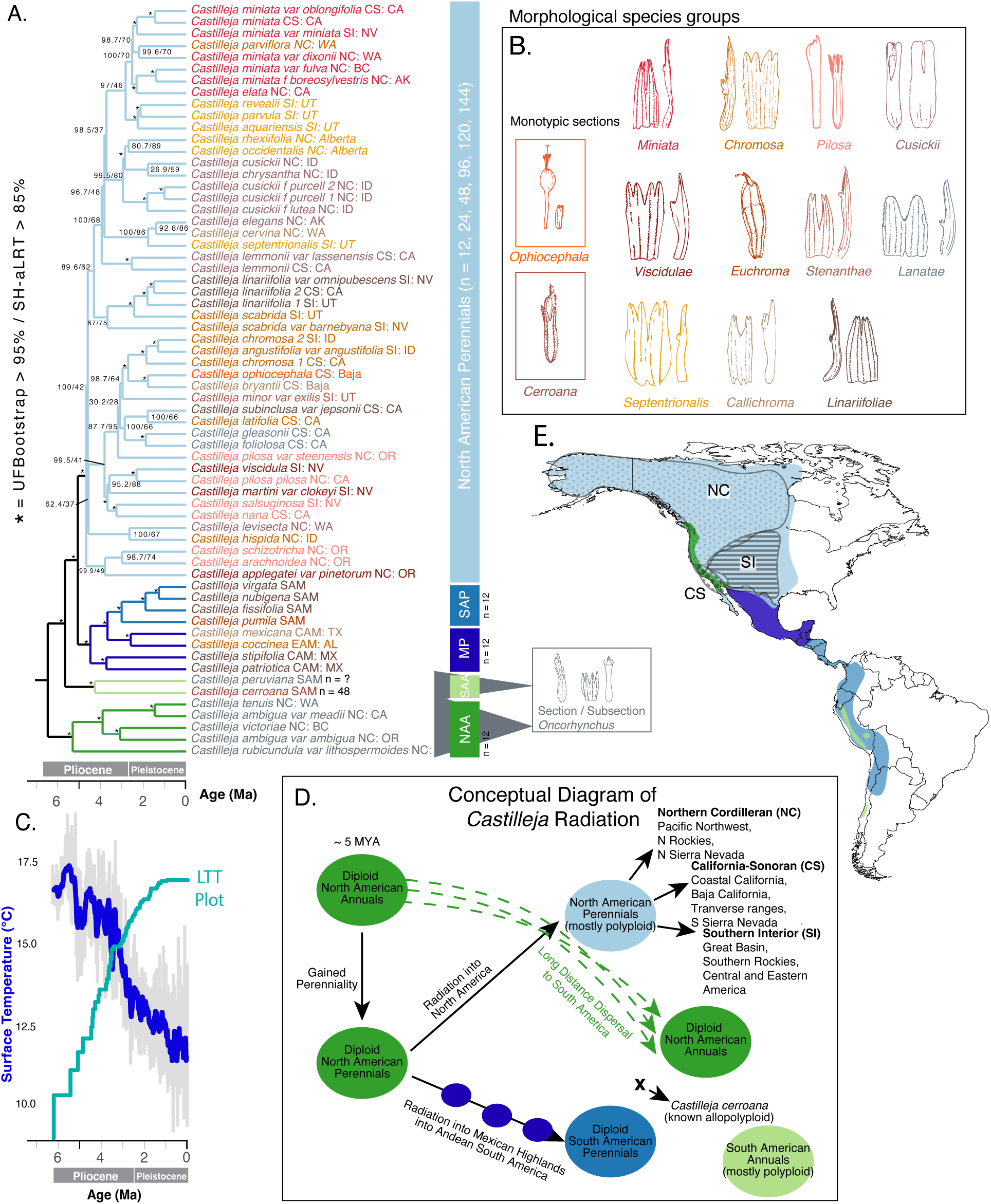
Integrative overview of *Castilleja* diversification, morphology, and biogeography: A) Concatenated dated phylogenetic hypothesis generated from filtered dataset with taxon names colors indicating morphological species groups shown in B and branches colored by geography and lifestyle (NAA = North American Annuals, SAA = South American Annuals, SAP = South American Perennials). Supported nodes (UFBootstrap > 95% and SH-aLRT > 85%) are depicted by stars at the nodes and unsupported nodes with listed support values. B) Morphological species groups showing corolla and calyx morphology designating species groups. C) Global Ocean surface temperature data over the last 7 Myr highlighting climatic fluctuations during time of *Castilleja* cladogenesis overlayed with a lineage through time plot of the phylogeny. D) Conceptual diagram of *Castilleja* radiation into North and South America with circles colored to indicate geography and lifestyle. E) Geographic ranges for the five clades in A with the three North American syngameon compartments shaded that correspond with chloroplast geographic structure.

Polyploidy in plants is often posited as a mechanism that promotes (nearly) immediate reproductive isolation, as whole genome duplication results in reduced fertility or hybrid infertility when polyploid individuals cross with either diploid relatives or those of differing ploidy levels (27–31). However, mechanisms including somatic doubling, the triploid bridge, and the hybrid bridge (32, 33) have been demonstrated to allow hybridization to occur across ploidy levels. In *Castilleja*, both auto- and allopolyploidy have been documented (18, 20, 21, 24–26, 34–38), yet these cytogenetic barriers do not appear to prevent gene flow entirely. Detailed morphological, molecular, and cytogenetic studies by Hersch-Green and Cronn (18) and Hersch-Green et al. (37) in a group of closely related Rocky Mountain *Castilleja* species have clearly demonstrated that different ploidy levels between species are only a weak barrier to hybridization, and that polyploidy and hybridization may play an important role in the generation of diversity in the clade.

In addition to biotic factors promoting hybridization, Plio-Pleistocene glacial oscillations may have facilitated historical hybridization in *Castilleja* by causing repeated cycles of expansion and contraction of suitable habitat resulting in concomitant rounds of geographic isolation and secondary contact.. While multiple studies have suggested that the perennial *Castilleja* clade has diverged relatively recently, within the last 3–5 mya (22, 23, 39), divergence time analyses with broad sampling have not yet been performed in the clade, making these hypotheses difficult to test. Taken together, these characteristics have led us to hypothesize that the recent and rapid radiation of *Castilleja* has resulted in weak reproductive barriers, thereby facilitating widespread introgression among species across the genus. Furthermore, given the topographic heterogeneity of the center of diversity of this clade (western North America), we propose that climatic fluctuations during the Plio-Pleistocene repeatedly created suitable habitat conditions that produced range contraction and expansion, leading to cycles of geographic isolation and connectivity between geographically restricted species pools or ‘compartments’. These alternating phases of isolation and secondary contact likely contributed to the formation of geographically structured syngameons within *Castilleja* (Figure 4). This pattern has also been suggested before in terms of ‘flickering connectivity’ seen in northern Andean páramos and other mountainous regions that experienced climatic changes during the Plio-Pleistocene (40–42).

Taxonomists have long struggled with clades of organisms that do not fit neatly into a bifurcating framework, as hypothesized morphological and ecological groups disintegrate when phylogenomic methods are applied. Even with increased genomic data (itself potentially discordant due to ILS, reticulate evolution or both), discordance between morphological and genetic patterns can persist. This study represents the first phylogenomic attempt to reconstruct the evolutionary history of this iconic clade of hemiparasitic wildflowers accounting for the fact that hybridization has always been hypothesized to contribute to the taxonomic and phylogenetic complexity of the clade (24, 25, 37). Using hundreds of newly sequenced nuclear genes for a geographically and morphologically representative sampling of *Castilleja*, we identify how hybridization has influenced its evolution. We provide evidence to support the clade as another important example of a syngameon—a phenomenon that is increasingly identified to explain the diversity and distribution of complex clades of both plants and animals, especially in heterogeneous regions of the world like western North America (1, 2, 4, 5, 11, 13). By carefully curating our phylogenomic dataset to control for paralogs and missing data, we construct a newly established phylogenetic hypothesis using concatenated, coalescent, and network-based methods to: (1) provide a temporal framework for exploring the broad biogeographic history of *Castilleja* and the factors that have promoted speciation, and 2) establish *Castilleja* as a syngameon by investigating gene tree discordance and hybridization in the context of the Edge-Range and Rapid Radiation hypotheses of syngameon generation and maintenance. We take the syngameon concept one step further by introducing the ‘multicompartmentalized syngameon’, a newly described pattern of historical and contemporary gene flow resulting from the dynamic climatic shifts of the Plio-Pleistocene. Whereas most documented syngameons involve relatively few interacting species within the same geographic area and similar ecological contexts, a multicompartmentalized syngameon encompasses numerous species interacting across broad spatial scales and heterogeneous environments. These interactions are organized into geographically and ecologically structured regions—here termed “compartments”—that broadly correspond to Level I ecoregions (43). We further suggest that multicompartmentalized syngameons may represent a common but previously underappreciated evolutionary outcome in species-rich, radiations shaped by repeated cycles of range expansion, isolation, and secondary contact driven by Plio-Pleistocene climatic dynamics.

## Results

### Sequence Recovery

After cleaning our raw dataset to remove adapter sequences and low quality reads, we recovered a mean of 1,009,448 reads per sample with a mean of 12% (1% - 25%) on target to the 353 nuclear loci (capture efficiency) using HybPiper2 (44). Since our sequence capture efficiency was low for some samples, we produced a filtered dataset to account for missing data. Our filtered dataset removed samples with less than 50% of genes and genes with less than 50% of taxa to produce a final dataset containing 262 genes (193,967 sites).

As expected, chloroplast by-catch recovered fewer reads than targeted nuclear data with a mean of 25,532 mapped reads per sample with a mean of 1.8% (0.5% - 8%) on target to the chloroplast coding sequences.

### Phylogenetic Relationships

#### Nuclear

Our nuclear concatenated analysis recovered four well supported clades corresponding to geography and life history (Figure 2A). Our ASTRAL analysis provided a similar but slightly different topology mainly corresponding to the placement of known allopolyploids (Supplementary Figure 1). However, ASTRAL can produce misleading topologies in the presence of hybridization (45), therefore we used the concatenated topology for our species tree hypothesis, while recognizing that any bifurcating model is biased when gene tree discordance is driven by introgression.

We recover a clade composed of North American annuals sister to all other *Castilleja* with high support (UFB = 100, SH-aLRT = 100) (Figure 2A). Within the remaining *Castilleja* we find South American annuals as sister to North and South American perennials. The South American annuals are composed of only *Castilleja cerroana* (a species of known allopolyploid origin) and *C. peruviana* (UFB = 100, SH-aLRT = 100) (21). Next, we recovered a well-supported clade composed of South American and Mexican perennials (UFB = 99.7, SH-aLRT = 99) with resolution towards the tips. Finally, we recovered a well-supported clade of North American perennials (UFB = 100, SH-aLRT = 100). Within the North American perennial clade, more recent nodes receive the highest support and we observe varying degrees of diminishing support with deeper nodes. Generally speaking, gene tree concordance with the species tree topology is highest at the deepest node of the tree, with minimal to moderate concordance within North and South American annual clades; in perennial clades, we find little to no concordance with the exception of a few shallow nodes (Figure 3A). The site concordance factors show that despite low or absent gene tree concordance, individual informative sites often retain moderate (25-50%) support for the species tree topology, while the remaining sites support alternative topologies, resulting in a discrepancy between site and gene concordance factors.

**Figure 3.**
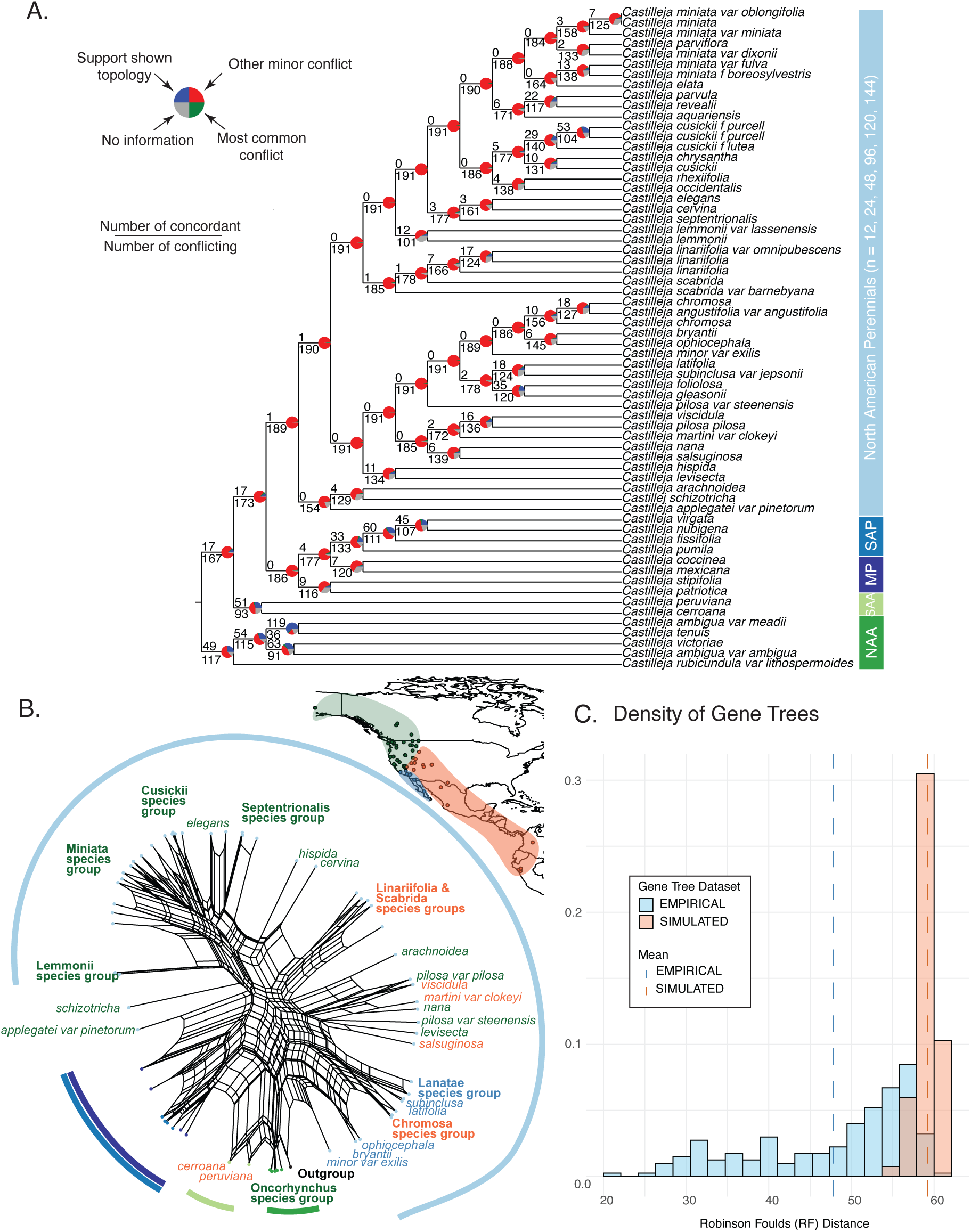
Gene-tree discordance patterns within our dataset. A) PhyParts analysis showing the number of gene trees that are concordant or in conflict with the concatenated phylogeny. Colored bars to left of phylogeny represent the geographic and lifestyle clades that are in Figure 2 (NAA = North American Annuals, SAA = South American Annuals, SAP = South American Perennials). B) Implicit species network derived from NANUQ distance matrix using SplitsTree revealing potential reticulation events and complex evolutionary relationships among taxa. Colored bars around tips correspond to colored bars in A. C) Coalescent simulations showing distribution of Robinson Foulds distance calculated from gene trees and ASTRAL-III tree as the empirical dataset compared to the distribution of the simulated dataset. The dotted line represents the mean of the distribution. The mean RF distance for the empirical and simulated dataset is shown with dotted lines.

#### Chloroplast

The chloroplast concatenated analysis resulted in three, low to moderately supported, strongly geographically structured clades (Figure 4). Within these clades, geographically proximate species tend to cluster together, while morphologically similar species are often not closely related.

**Figure 4.**
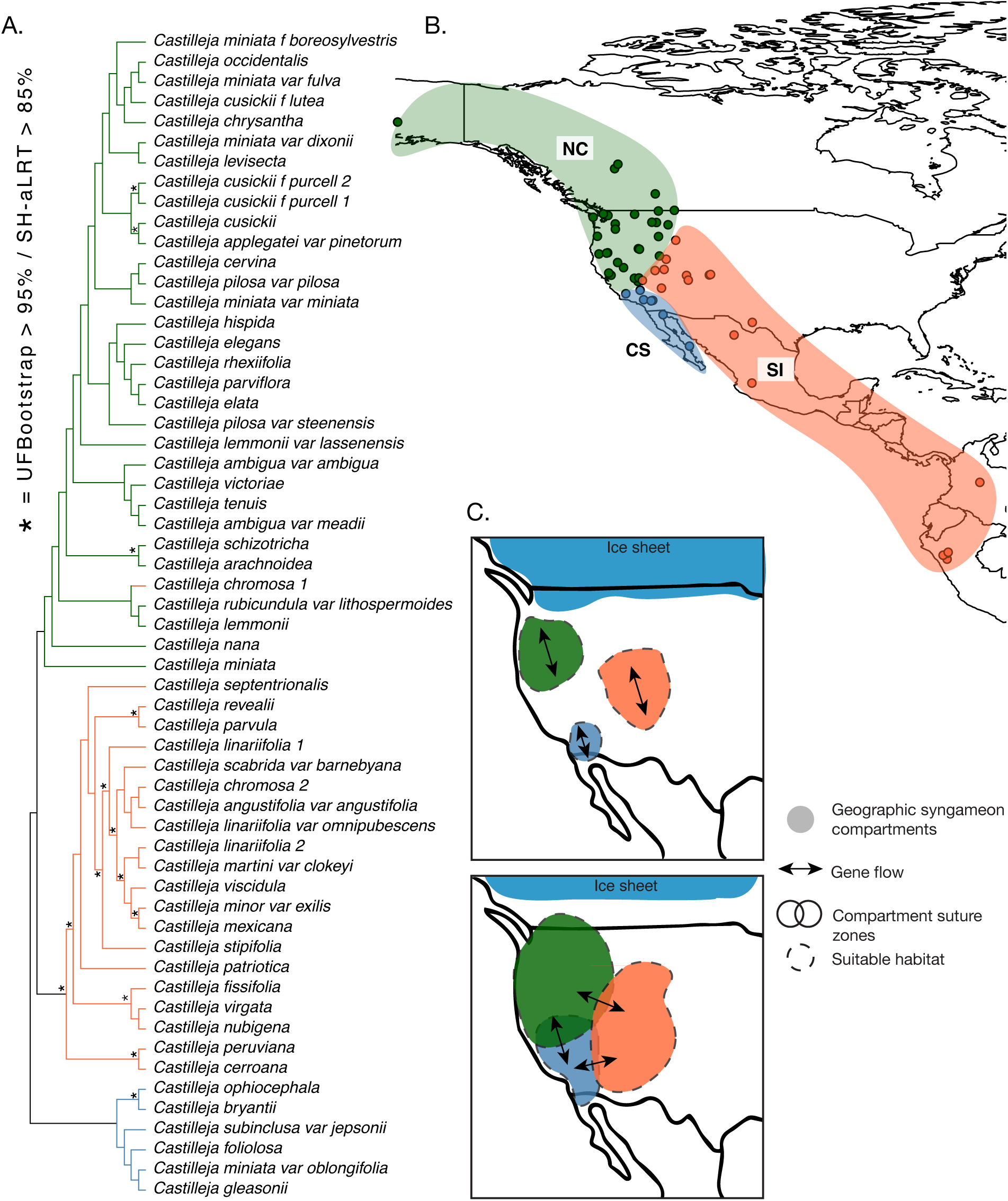
A) Concatenated chloroplast phylogenetic hypothesis with branches colored with respect to their geographic area in B. Supported nodes (UFBootstrap > 95% and SH-aLRT > 85%) are depicted by stars at the nodes. B) Biogeographic distribution of major clades corresponding to the bifurcating chloroplast topology. C) Conceptual diagram illustrating the contraction and expansion of syngameon compartments in response to Plio-Pleistocene climatic fluctuations. During warm interglacial periods, species ranges expand and overlap, facilitating widespread gene flow and the formation of interconnected syngameon networks via suture zones. In contrast, glacial periods drive habitat fragmentation and range contractions, isolating populations into distinct compartments localized hybridization within the syngameon compartment. These cyclical climatic shifts repeatedly reconfigure the structure and connectivity of syngameons over time.

### Coalescent Simulations

The distribution of tree-to-tree distances from the empirical gene trees to the species tree and simulated gene trees to the species tree is broadly non-overlapping (40% overlap), indicating a low degree of similarity between the empirical and simulated distributions (Figure 3C). Therefore, we infer that only a portion of the gene-species tree discordance can be attributed to coalescent stochasticity under an ILS-only simulation. RF distances to the species tree are lower from the empirical gene trees than the simulated gene trees. This may be due to the simulated gene trees being generated under the ASTRAL topology and then compared to the concatenated species tree, model differences between the tree that genes were simulated under and the tree that they were compared to could lead to increased RF distances.

### Hybridization Analyses

We used NANUQ to summarize gene tree discordance and visualize potential reticulate relationships via gene quartet distributions constructed from 298 gene trees. The resulting network closely mirrors well-supported clades in the concatenated species tree topology but reconstructs one main cluster of all *Castilleja* indicating potential hybridization across the major morphological and geographic clades (Figure 3B). One dense cluster of cycles encompasses the South American perennial clade, including *Castilleja peruviana* and *C. cerroana*, with edges connecting these taxa to the North American annuals and South American perennials, consistent with historical introgression previously documented between these geographically disjunct lineages (21) (Figure 3B). Another prominent cluster of cycles is within the North American perennials representing the Miniata, Septentrionalis, and Cusickii species groups (Figure 2B, 3B). Other, less prominent clusters of cycles include most of the hypothesized morphological species groups seen in the bifurcating topology such as the Pilosa, Linariifoliae, Chromosa species groups (Figure 2B, 3B). The presence of these cycles suggests that gene tree discordance in these regions may result from hybridization, warranting further investigation with explicit network models.

### Divergence Dating

The dated phylogenetic tree estimated the age of the most recent common ancestor (MRCA) of *Castilleja* is 6.4 Myr (95% CI: 5.2-7.6) and we observe that most of the cladogenesis events have taken place in the last 2-6 Myr (Figure 2A, 2C, Supplementary Figure 2). The MRCA of the Central American perennial grade and North American perennial clade was dated at 5 Myr (95% CI: 4-6).

## Discussion

Hybridization is increasingly recognized as an important feature of recent evolutionary radiations, but its consequences are typically studied among small numbers of closely related, geographically overlapping species (20, 21, 46–50). Our phylogenomic and network-based analyses reveal a different pattern in *Castilleja*: extensive gene flow among non-sister lineages distributed across much of North America. We detected recent introgression primarily within geographic regions and widespread historical introgression among regions, indicating that the evolutionary history of this clade of ∼200 species cannot be adequately represented by a strictly bifurcating phylogeny. Instead, *Castilleja* forms a geographically structured network of interacting lineages that we term a *multicompartmentalized syngameon* (MCS). This framework helps explain both the persistent difficulty of reconstructing relationships in the clade (20, 21) and how extensive gene flow can remain spatially structured across a continental radiation.

The temporal history of *Castilleja* provides a mechanism for the origin of this structure. We estimate the crown age of the genus at ∼6.4 Myr and the onset of the perennial clade at ∼5 Myr, with most extant diversity emerging during the Plio-Plestocene (Figure 2C and 2D). Rapid diversification over this interval likely limited the evolution of strong reproductive isolating barriers, allowing repeated hybridization to contribute to gene-tree discordance observed across the nuclear genome. At the same time, Plio-Pleistocene climatic oscillations repeatedly contracted, fragmented, and expanded suitable habitats across western North America (40–42). These changes would have alternately isolated lineages and brought them into secondary contact, promoting recurrent gene flow within geographic compartments and episodic exchange among compartments. The MCS therefore reflects the joint effects of rapid radiation and repeated, climate-driven changes in geographic connectivity.

The contrast between the chloroplast and nuclear phylogenies provides a genomic record of these dynamics. The chloroplast DNA (cpDNA) phylogeny strongly groups lineages by geography rather than by morphological species relationships (Figure 2B), and most clearly identifies the geographic compartments (Figure 4). Because chloroplast genomes are predominantly maternally inherited, hybridization followed by repeated backcrossing can retain a locally acquired chloroplast genome while largely retaining the nuclear genome of the recurrent parent, a process known as chloroplast capture (51–53). Repeated chloroplast capture among co-occurring lineages can thus cause cpDNA relationships to track geography more closely than species history. In contrast, the nuclear phylogeny more closely recovers morphologically defined groups while retaining extensive conflict produced by historical hybridization and lineage connectivity within and among compartments (Figures 2 and 3). Together, these complementary signals reveal a system in which species identities persist even as different components of their genomes record different dimensions of a shared reticulate history. Although incomplete lineage sorting of organellar haplotypes, restricted seed dispersal relative to pollen dispersal (54), genetic drift associated with population bottlenecks and serial founder events during range contractions and expansions (55–57), and selection on chloroplast haplotypes or cytonuclear combinations (58) could also contribute to geographic cpDNA structure. The marked discordance with the nuclear phylogeny (Figures 2A, 4A), together with independent evidence of widespread introgression (Figure 3B), is consistent with repeated chloroplast capture as an important mechanism.

### The multicompartmentalized syngameon (MCS) framework

Classic syngameons comprise morphologically distinct species connected by ongoing or episodic gene flow. Selection can maintain species differences despite hybridization and backcrossing, while introgressed variation may facilitate adaptation and expansion into new ecological settings (6, 13, 16, 59, 60). Two major hypotheses describe how these systems arise and persist. The rapid radiation hypothesis emphasizes incomplete reproductive isolation during short intervals of diversification, whereas the edge-range hypothesis emphasizes hybridization when shifting range margins bring species into contact, potentially allowing beneficial, locally adapted alleles to cross species boundaries (13, 16). The *Castilleja* MCS unites these two processes across a broader spatial and temporal scale: a rapid radiation partitioned into multiple, semi-discrete geographic compartments whose connectivity changes through time. Conceptually, this dynamic resembles fusion and fission documented in African cichlid radiations and cycles of isolation and migration in mangroves, but extends those processes to a persistent network of multiple regional species assemblages distributed across continental scales (5, 40).

We identify three broad geographic compartments—the Californian-Sonoran (CS), Northern Cordilleran (NS), and Southern Interior (SI)—from the spatial clustering of lineages in the cpDNA phylogeny rather than from predefined ecoregion boundaries (Figure 4). Figures 2E and 4 provide complementary views of this structure: Figure 2E summarizes the broader distributions of the major geographic lineages recovered from the nuclear phylogeny, whereas Figure 4 shows the sampled localities and geographic compartments recovered from cpDNA analyses. The Californian-Sonoran compartment (CS; Figure 4B) spans the transition between the Mediterranean-climate flora of southern California and northwestern Baja California and the seasonally arid floras of the Baja Peninsula and adjacent southwestern deserts. It is centered on the southern California coast and extends through the Transverse Ranges and southern Sierra Nevada into Baja California and adjacent portions of the southwestern deserts, lying primarily within the Mediterranean California and North American Deserts Level I ecoregions (43). The Northern Cordilleran compartment (NC; Figure 4B) extends from the Coast Ranges of California and central Sierra Nevada northward through the Pacific Northwest to Alaska and eastward into the northern and central Rocky Mountains. It encompasses primarily the Marine West Coast Forests and Northwestern Forested Mountains ecoregions, with northern extensions into the Taiga and Tundra. The Southern Interior compartment (SI; Figure 4B) is centered on the Great Basin and southern Rocky Mountains but extends southward through the southwestern United States and Texas into Mexico and, more broadly, Central and South America (Figure 2E). This compartment also includes a small number of *Castilleja* species occurring in the Great Plains and eastern North America. Within Canada, the United States, and Mexico, this SI compartment crosses portions of the North American Deserts, Southern Semi-Arid Highlands, Temperate Sierras, Tropical Dry Forests, Great Plains, and Eastern Temperate Forests. Thus, the compartments are broad geographic entities that contain mosaics of Level I ecoregions rather than units defined by, or uniquely corresponding to, individual ecoregions. They are neither discrete clades nor fixed biogeographic units, but probable domains of evolutionary interaction; gene flow occurs more frequently within them than among them, while their boundaries shift and remain permeable through time.

We hypothesize that Plio-Pleistocene climatic oscillations repeatedly altered the distribution and connectivity of suitable habitat within these broad geographic compartments (Figure 4C). During glacial phases, suitable environments may have become spatially restricted or regionally aggregated, concentrating *Castilleja* lineages within shared refugial areas and thereby increasing opportunities for hybridization within compartments. This mechanism has an empirical precedent in the angiosperm clade Heuchereae (Saxifragaceae), where most chloroplast capture events date to the mid- to late-Pleistocene and ancestral niche reconstruction indicates that climate-driven range contraction brought previously allopatric lineages into contact within shared refugia (51). Subsequent interglacial expansion would have redistributed these lineages across broader geographic areas while preserving the genomic consequences of gene flow that occurred within regional refugia. Because expansion generally proceeded from separate regional source areas, contact remained more frequent among lineages belonging to the same compartment than among lineages from different compartments. Gene flow among compartments would have occurred less frequently, primarily where expanding ranges met in geographic transition or suture zones. This model provides a potential mechanism for the combination of frequent recent introgression within compartments and less frequent, predominately historical connectivity among them observed in our analyses. It predicts more frequent and recent introgression within compartments, geographically clustered organellar capture, and less frequent, older introgression among compartments concentrated near historical suture zones.

These cyclical changes in habitat connectivity are expected to leave different but complementary signatures across genomes and spatial scales. Hybridization and backcrossing within regional refugia promote the geographic sorting and capture of uniparentally inherited genomes, causing organellar relationships to become more strongly associated with geography than with morphology. In contrast, nuclear genomes retain evidence of broader historical relationships while more closely tracking morphological species relationships. Thus, the defining feature of an MCS is not simply extensive hybridization or geographic structure, but their interaction through time: recurrent gene flow within regional species networks combined with intermittent connectivity among those networks. Episodic contact among compartments may also allow alleles acquired within one regional species network to propagate through the broader syngameon, providing a potential route for adaptive variation to move across otherwise differentiated geographic and ecological contexts (2, 4, 6, 11, 13, 16). Such an adaptive role for interspecific gene flow is supported by genomic evidence that hybridization contributes to parallel adaptation in *Vitis* (61). Identifying the genomic regions that cross (or resist) compartment boundaries will be necessary to test this possibility. Although we do not reconstruct the locations or extent of suitable habitat during individual glacial cycles, the spatial and temporal distribution of introgression observed here motivates this refugial-concentration hypothesis and generates testable predictions for future paleoclimatic and population-genomic analyses.

Several features of our data support hybridization, rather than incomplete lineage sorting alone, as the primary source of this reticulate pattern. The North American perennial backbone is poorly resolved, and nuclear gene trees show substantial conflict (Figure 3A), yet the discordance expected from incomplete lineage sorting only accounts for a small proportion of the total observed conflict (Figure 3C). Network analyses instead identify widespread introgression across the clade (Figure 3B), consistent with the contrast between geographically structured cpDNA lineages and the greater morphological cohesion recovered from nuclear data. Denser population-level sampling will be required to estimate the relative timing and direction of individual introgression events, but the combined evidence indicates that neither a single species tree nor incomplete lineage sorting alone can explain the genomic history of the group.

Polyploidy, although prevalent in perennial *Castilleja*, has also not imposed a complete barrier to gene flow. Experimental crosses among co-occurring octaploid *C. miniata*, tetraploid *C. rhexiifolia*, and diploid *C. septentrionalis* resulted in observed gene flow across both species and ploidy boundaries, although barriers were weaker in conspecific crosses and between individuals with compatible chromosome complements (37). Polyploidy may therefore shape the probability and direction of introgression without severing the connections that maintain the MCS. More broadly, this result cautions against treating cytogenetic differences as sufficient evidence that rapidly radiating lineages are evolutionarily independent.

The MCS framework changes the goal of phylogenetic reconstruction in *Castilleja*. Rather than seeking a single bifurcating topology that eliminates conflict, future work should characterize how evolutionary histories vary across the genome and landscape. Denser geographic and population-level sampling— particularly of organellar variation within widespread species and across complete regional species assemblages—will be necessary not only to delimit compartments more precisely, identify gradients of compartment membership, and locate transition zones and potential suture zones (Figure 4), but also to identify additional climate-driven evolutionary dynamics within compartments that cannot be resolved from the present sampling, including peripatric divergence and incomplete speciation associated with recurrent range fragmentation (62). Comparable nuclear sampling can then determine which genomic regions move across, or resist, compartment boundaries. Coupling these data with paleodistribution and ancestral niche models offers a direct way to test whether present-day compartment boundaries reflect historically recurrent zones of habitat overlap and hybridization (51). Because introgression may involve multiple co-occurring species, sampling only from focal species pairs or morphological complexes risks omitting important routes or contributors of gene flow, thereby misrepresenting the broader evolutionary network. Similar multicompartmentalized syngameons may occur in other species-rich radiations shaped by repeated climatic oscillations, particularly where topographic and/or ecological heterogeneity alternately isolates regional biotas and brings them back into contact. Recognizing this structure provides a way to move beyond the choice between geographic history and species history; in reticulate radiations, both are preserved, but in different genomes and at different spatial scales.

## Material and Methods

### Taxon Sampling

We included 70 accessions representing 55 of the 217 currently recognized species of *Castilleja* with a focus on North American perennial species. This sampling was selected to broadly represent the morphological and biogeographical sections proposed by J.M. Egger, D.C. Tank, and S.J. Jacobs based on their 80+ combined years of collections-based research in the clade (unpubl.data); we also included a small sampling of intraspecific taxa for several widespread species. We sampled two outgroup taxa in the closely related genera *Chloropyron* and *Cordylanthus* (11) (Supplementary Table 1). Leaf tissue samples were obtained from field collections and dried using silica gel desiccant with corresponding vouchered herbarium collections curated at ID, WTU (Supplementary Table 1).

### Molecular Protocols

#### DNA Extraction

DNA was extracted from 20-30 mg leaf tissue using a modified CTAB method (63). Leaf tissue was ground using a Mixer Mill MM400 (Retsch, GmbH, Haan, Germany). We incubated the ground tissue with a 2% CTAB buffer overnight at 65°C and precipitated it in isopropanol at 2°C for 30 minutes. DNA was quantified by a Qubit fluorometer (Thermo Fisher Scientific, Inchinnan, UK) and visualized by running a 1% agarose gel to assess fragment distribution. Extracted DNA was cleaned with a 1.6X Sera-Mag SpeedBeads (GE Healthcare Life Sciences, Pittsburgh, PA, USA) to remove any inhibitory impurities and very small fragments.

#### Library Preparation

All input DNA was normalized to 10 ng/uL for library preparation. Library preparation was completed using Illumina’s Nextera tagmentase approach (Illumina, San Diego, CA, USA) at ¼ volumes to reduce per sample cost. We re-amplified using KAPA Hifi kit (13 cycles; 1:00 elongation). Prepped libraries were initially cleaned with 1.6X MagBio beads (MagBio Genomics, Gaithersburg, MD, USA) to remove primers. We then performed a dual-sided size selection with MagBio beads to select fragments within our desired range for the MiSeq (360 – 1200bp). For our pilot study, we pooled 12 samples with 20-25ng of cleaned library. For the other samples, we pooled 8-12 samples with 25-60ng of cleaned library. We analyzed fragment distribution by running pooled samples on 5200 Fragment Analyzer System (Agilent, Santa Clara, CA, USA) to confirm they were within our desired distribution before moving on with capture.

#### Target Capture

We used the Angiosperm-353 myBaits^®^ Expert Panel target capture kit (Arbor Bioscience, Ann Arbor, MI, USA). We ran pooled libraries through target capture at quarter reactions to decrease cost. Pools were hybridized at 65°C for 24 hrs and then re-amplified with KAPA HiFi HotStart ReadyMix kit (25 cycles and 1:00 elongation) (KAPA BioSystems, Cape Town, South Africa). Following re-amplification, pooled libraries were cleaned with MagBio beads to remove the primers. To confirm target capture worked we quantified the amount of pooled library with a Qubit fluorometer using a broad range, double stranded assay kit (Thermo Fisher). Pools were then combined into a final pool for sequencing. For known polyploids we doubled the amount of sample for the final pool to be able to phase polyploids (Supplementary Table 1).

#### Sequencing

Sequencing was performed on an Illumina MiSeq v3 (600 cycles of 2 x 300 bp paired-end reads) chemistry (Illumina, San Diego, CA) at the IIDS Genomics and Bioinformatics Resources Core at the University of Idaho and on an Illumina NextSeq v1000 (P1 chemistry: 600 cycles 2 x 300 bp paired-end reads) at the Integrated Microscopy Core at the University of Wyoming. Sequencing proceeded in three batches, resulting in 65 accessions of 50 species.

### Data Preparation

#### Read Mapping

We first cleaned raw reads with Trimmomatic to remove adaptor sequences and low-quality bases (64). Cleaned reads were mapped to the Angiosperm-353 nuclear target genes and coding regions of the chloroplast using HybPiper v2.3.2 (44). To maximize nuclear sequence recovery, we used NewTargets to create a new, more specific target file (65) tailored to Orobanchaceae taxa (66). Read mapping was performed using HybPiper v2.3.2, which employs the Burrows–Wheeler Aligner (BWA) for initial read recruitment to target sequences (67). Gene recovery and assembly were conducted using the HybPiper v2.3.2 workflow, which integrates reference-guided assembly approaches. Exon sequences were aligned and scaffolded using *exonerate*, while intronic regions were recovered using *run-intronerate* to extract non-exonic sequences (68). Finally, we used *retrieve sequences* to compile both exon and intron for downstream phylogenomic analyses.

A similar process was used for recovering chloroplast data from coding sequences. The chloroplast target file was made by retrieving all coding regions from the previously sequenced *Castilleja paramensis,* which amounted to 50 genes (Genbank ID NC_031805). Raw reads were mapped to targets using HybPiper v2.3.2 with the same parameters from the nuclear assembly.

#### Sequence Alignment and Filtering

After recovering nuclear supercontig sequences from HybPiper v2.3.2, nuclear sequences were aligned using MAFFT (69). We used ParaGone v1.1.3 to resolve paralogous sequences and infer orthology using algorithms described in Yang and Smith 2014 and implemented in Morales-Briones 2022. We used the monophyletic outgroup (MO) algorithm as our tree-based decomposition approach. This method retains only gene trees with a monophyletic outgroup, roots the tree accordingly, and iteratively identifies and prunes duplication. After inferring orthology, we evaluated the impact of missing data using a custom python script (https://github.com/santosmalia/Castilleja_Angio353). The script filtered out samples if they are not present in a percentage of the genes (we examined 50%) and subsequently removed genes lacking representation in at least (50%) of the samples, hereafter referred to as the filtered dataset.

Chloroplast sequences were aligned with MAFFT, cleaned with phyutility to remove sites that are not present in at least 30% of taxa (69, 70). We manually inspected and edited alignments in AliView to remove spurious alignments due to low coverage or missing data in some taxa. Further, we removed taxa with low coverage.

### Phylogenetic Tree Reconstruction

#### Nuclear Gene Trees

We generated initial gene trees for each filtered nuclear dataset in IQ-TREE2 v2.1.1 with 1000 ultrafast bootstrap (UFB) replicates and 1000 regular bootstrap replicates (BS) (71–73). ModelFinderPlus was used to choose the appropriate model of sequence evolution (74).

#### Nuclear Species Tree

##### Concatenation

We implemented a partitioned analysis with model testing in IQ-TREE2 v2.1.1 for each filtered dataset using PartitionFinder (-p, TESTMERGE) with support using 1000 ultrafast bootstrap replicates and the Shimodaira-Hasegawa-like approximate likelihood ratio test (SH-aLRT) (75) to assess nodal support (Figure 2A). To visualize gene tree discordance we used the open source software PhyParts (https://bitbucket.org/blackrim/phyparts/src/master/) to calculate the proportion of gene trees that were concordant with, in conflict with, and uninformative for each node of the species tree (76).

##### ASTRAL-III

We used ASTRAL-III to estimate a species tree from gene trees generated as described above. ASTRAL-III has been shown to be statistically consistent under the multi-species coalescent model (77) and produces a topology accounting for incomplete lineage sorting (ILS). We generated branch support for the topology with local posterior probabilities (LPP) and quartet support (QS). LPP is the probability that a branch is the true branch given the set of input gene trees, where QS is the proportion of quartets (unrooted 4-taxon tree) in the gene trees that support the given branch. We also produced the QS for the two alternative topologies at each node in ASTRAL-III using the -t 2 flag. We employed the polytomy test available in ASTRAL-III to test that the null hypothesis that a branch is a polytomy can be rejected (-t 10 flag).

#### Plastid Tree

We generated a concatenated analysis in IQ-TREE2 v2.1.1 for the final dataset using General Time Reversible (GTR) model of substitution with support using 1000 ultrafast bootstrap replicates and the Shimodaira-Hasegawa-like approximate likelihood ratio test (SH-aLRT) (75) to assess nodal support.

### Hybridization Analyses

To understand the extent of hybridization within *Castilleja* we conducted a broad analysis of hybridization by constructing a split network based on gene trees produced above and quartet likelihood using the method Network inference Algorithm via NeighborNet Using Quartet distances (NANUQ) in MSCquartet R package, v. 2.0 (78, 79). NANUQ aims to provide estimation of level-1 species networks (bifurcations and reticulation events), which takes gene trees and calculates gene quartet distributions under the Network Multispecies Coalescent model. NANUQ implements a two-hypothesis testing framework defined by parameters: ɑ and ꞵ, where smaller values of ɑ and ꞵ indicate more conservative thresholds for inferring tree-like versus reticulate relationships. We explored a range of ɑ values as suggested by authors and chose values based on visual inspections of simplex plots, to assess whether the quartets exhibit a signal of network cycles. The resulting distance matrix was then used to create a splits graph under the Neighbor- Net algorithm in the SplitTree App (80, 81).

### Coalescent Simulations

To understand if ILS alone can explain the gene tree discordance, we used a simulation based approach following previous literature (82, 83). We first constructed an ultrametric species tree with branch lengths in coalescent units (T/4N_e_), hereafter referred to as our constrained tree, by using our ASTRAL-III topology to perform a constrained tree search in IQ-Tree2 under a strict molecular clock, producing a constrained tree in mutational units (uT). We estimated the population size parameter, θ from the mutational branch lengths in the constrained tree and the coalescent branch lengths from the ASTRAL-III tree for internal branches. θ was set to one for terminal branches. We simulated 10,000 gene trees based on our constrained tree using the R package Phybase v.2.0 under a multi-species coalescence (MSC) model from Rannala and Yang (2003) (70, 84). After generating simulated gene trees, we used phyutility to root the trees on the appropriate outgroup (85). Using TreeCmp, we then calculated Robinson-Foulds (RF) tree-to-tree distances of simulated gene trees and the empirical species tree (39). We then compared this value to the RF distance between our empirical gene trees and the empirical species tree to quantify how much ILS was contributing to gene tree discordance. The ratio of the mean RF distances between the simulated and empirical datasets was calculated as a measure of the amount of observed gene-tree discordance caused by ILS.

### Divergence Dating

We used BEASTv2.7.7 to conduct the divergence dating analysis. Due to the lack of fossils in Orobanchaceae, we used a secondary calibration approach where we matched concordant nodes of our phylogeny from previously published phylogenies (86). We added 10 additional outgroup taxa from publicly available databases to place secondary calibrations on outgroup nodes (Supplementary Table 1) (39). The phylogenetic positions of outgroup taxa were constrained to the relationship estimated by Schneider and Moore (2017) (39). Due to the nature of missing data in our dataset, we used the top 10 genes with the most coverage in BEAST v2.7.7 to estimate divergence dates. We enforced a fixed topology in our BEAST analysis. To retrieve a topology with appropriate outgroups for placing calibrations, we fixed outgroup relationships to match from Schneider and Moore (2017). We fixed ingroup relationships to match relationships inferred from the concatenated analysis of our dataset with paralogs resolved. We used AMAS to concatenate the final 10 genes and create a nexus to input to Beauti (87). We ran two independent analyses under the optimized relaxed clock model and the Yule model for 50 million generations. We combined runs using LogCombiner v2.7.7 with a burnin of 10%. Convergence and ESS values were checked in Tracer v1.7.2 and the posterior sample of phylogenetic trees was summarized as a maximum clade credibility tree using TreeAnnotator v2.7.7. We also completed a run with no data and sampling from the prior to assess goodness of fit. A lineage through time plot (LTT) was also created using phytools (88) to visualize cladogenesis events during major climatic events (89) (Figure 2C).

## Acknowledgments

Thank you to Dr. Jack Sullivan, Dr. Marjorie Weber, Dr. Luke Harmon, Dr. Catherine Wagner, Dr. Melanie Murphy, and Dr. Matt Carling for comments and discussion that improved early versions of this paper. We are also grateful to the Associate Editor and anonymous reviewers for their valuable feedback. S.H. was supported by an Institutional Development Award (IDeA) from the National Institute of General Medical Sciences of the National Institutes of Health under grant number 2P20GM103432.

## Data Availability Statement

The raw sequencing reads generated in this study have been deposited in the GenBank Short Read Archive (SRA) under accession number XXXXX. All scripts used for data processing and analysis are available on GitHub at https://github.com/santosmalia/Castilleja_Angio353.

---

**Table 1.** Secondary calibration priors taken from Schneider and Moore (2017) used to generate a time-calibrated tree for *Castilleja* and related taxa in BEAST. We used lognormal priors in the BEAST analysis. Mean ages are shown for calibration points along with the reported 95% confidence intervals.

| Lineage | Schneider et al 2017 |  | This study |  |
| --- | --- | --- | --- | --- |
|  | Crown Age (Ma) | 95% HPD (Ma) | Crown Age (Ma) | 95% HPD (Ma) |
| Orobanchaceae (including Rehmanniaceae) | 30.2 | 25.6-36.0 | 27.1 | 23.20-31.0 |
| Pedicularideae + Buchnereae + Rhinanthaeae | 24.1 | 20.4-28.9 | 21.7 | 18.7-24.7 |
| *Castilleja* + *Triphysaria* | 5.0 | 3.8-6.2 | 7.6 | 6.3-8.9 |
| *Castilleja* | - | - | 6.4 | 5.2-7.6 |

## Supplemental Figures & Tables

**Supplemental Figure 1.** Coalescent species tree inferred by ASTRAL-III, depicting our best hypothesis for relationships within *Tricalysia* and closely related genera compared to concatenated IQTree2 topology.

**Supplemental Figure 2.** Divergence dating analysis results with secondary calibrations from Schneider and Moore (2017).

**Supplemental Table 1.** List of *Castilleja* samples and outgroups used in this study with associated collector name and numbers, highlighting the morphological sections, biogeographic regions, perenniality, and known ploidy estimates.

## Notes

### Competing Interest Statement

The authors have declared no competing interest.

