## Supplementary figures and images for "Painting the evolutionary history of a multicompartmentalized syngameon in North American *Castilleja* (Orobanchaceae)"

### Supplemental Figure 1

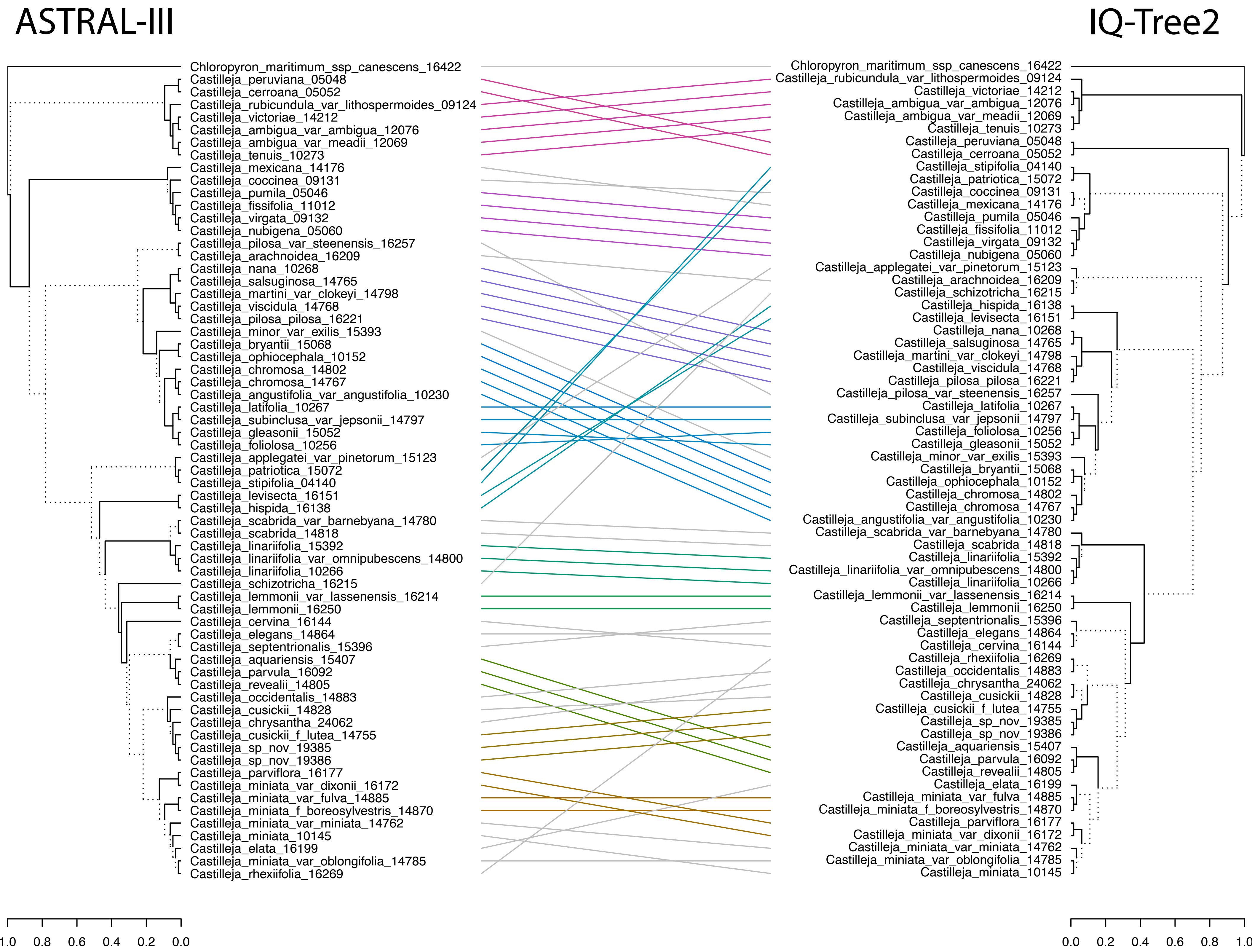

### Supplemental Figure 2

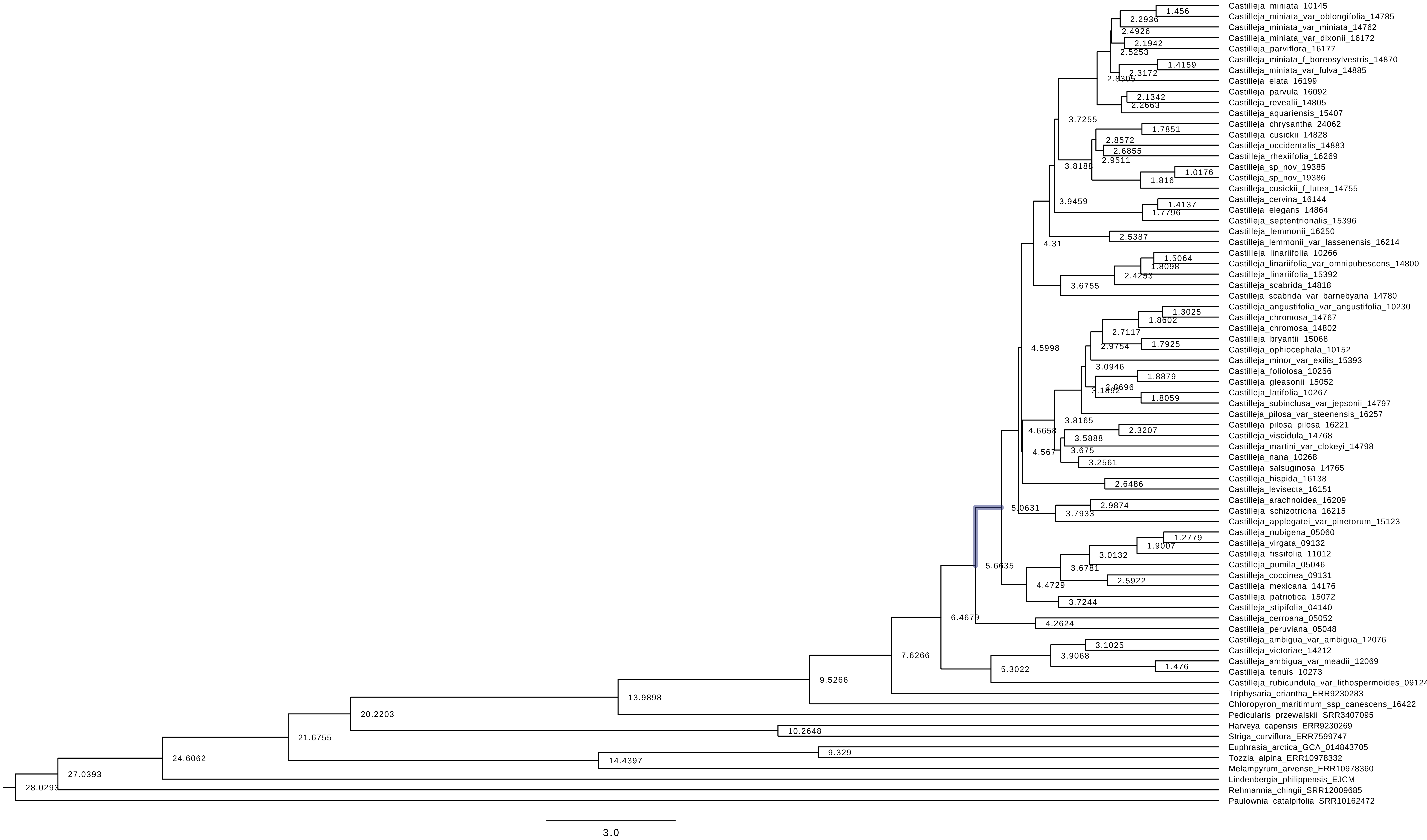
